# ya||a: GPU-powered Spheroid Models for Mesenchyme and Epithelium

**DOI:** 10.1101/525352

**Authors:** Philipp Germann, Miquel Marin-Riera, James Sharpe

## Abstract

ya||a is yet another parallel agent-based model for morphogenesis. It is several orders of magnitude faster than conventional models, because it runs on GPUs and because it has been designed for performance: Previously only complex and therefore computationally expensive models could simulate both mesenchyme and epithelium. We chose to extend the simple spheroid model by the addition of spin-like polarities to simulate epithelial sheets and tissue polarity. We also incorporate recently developed models for protrusions and migration. ya||a is written in concise, plain UDA/C++ and available at github.com/germannp/yalla under the MIT license.

## Introduction

Embryonic tissues come in two basic states, epithelial and mesenchymal. Epithelial cells typically create compact tissues with strong intercellular contacts, thus acting as physical barriers to other cells and molecules. Epithelial tissues are organized into sheets with stable neighbourhoods and show a marked apical-basal polarity across the depth of the sheet. Such sheets undergo complex 3D deformations by means of active cell behaviors, which result in folding, bending, or twisting of the sheet, or by means of passive mechanical forces exerted by the surrounding tissues (Honda, 2017). Mesenchymal cells, on the other hand, are typically looser 3D tissues with abundant extracellular matrix. Mesenchymal cells come in various shapes which are typically highly dynamic due to the formation and retraction of protrusions (such as filopodia, lamelopodia, etc.). Mesenchymal tissues change shape by proliferation, extracellular matrix secretion and remodeling, and intercalation (G. W. Brodland and H. H. Chen, 2000). Cell transitions from epithelial to mesenchymal type, and interactions between both tissue types, are common throughout development and disease.

Previous simulation frameworks for morphogenesis (Hoehme and Drasdo, 2010; Richmond et al., 2010; Gorochowski et al., 2012; Rudge et al., 2012; Swat et al., 2012; Mirams et al., 2013; Sütterlin et al., 2013; Kang et al., 2014; Starruß et al., 2014; Cytowski and Szymanska, 2015; Barton et al., 2017; Somogyi and Glazier, 2017; Sussman, 2017; Ghaffarizadeh et al., 2018; Song et al., 2018) often emphasized either epithelial or mesenchymal processes, e.g. vertex models describe the shapes of epithelial cells within sheets or cellular Potts models describe differential adhesion (Osborne et al., 2017). Recently, solutions to overcome this limitation were put forward: an extension of the spheroid model by torsion joints (Disset et al., 2015), a 3D implementation of the vertex model (Okuda et al., 2015), a sub-cellular element model using apical and basal elements for epithelial cells (Gord et al., 2014), a sub-cellular element model using cylindrical elements for epithelial cells (Marin-Riera et al., 2015), and most recently a spheroid model with apical-basal polarity (Delile, Herrmann, et al., 2017).

However, none of these frameworks natively supports all the diverse mesenchymal and epithelial cellular behaviors and all are computationally more complex than necessary. We therefore chose to write a new simulator, dedicated for running on graphics processing units (GPUs) to take advantage of their highly efficient parallelized speed. Our new simulator extends the classical spheroid model by incorporating concepts from magnetism to simulate the apical-basal polarity of epithelia and the tissue polarity seen in mesenchyma. We also added an implementation of recent methods to model contractile protrusions (Belmonte et al., 2016; Palsson and Othmer, 2000) and individual cell migration (Delile, Doursat, et al., 2014).

## Results

In a spheroid model a cell *i* is described by the center of its spheroid 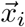. Neighboring cells interact via spherical potentials with a repelling core and an attractive zone around it (Figure 1 A). Cells in contact can exert friction on each other (Okuda et al., 2015) and can exchange chemical signals (Figure 1 A, B). To enable convergent-extension and efficient cell sorting we added contractile cellular protrusions, which allow more distant cells to pull on each other (Belmonte et al., 2016; Palsson and Othmer, 2000) (dotted line in Figure 1 A). These links are created and destroyed repeatedly over time and when a field of such links is oriented in a non-random fashion it leads to convergent-extension, as demonstrated in Belmonte et al. (2016) and in Figure 1 C. Alternatively, if the contractile protrusions are randomly oriented but preferentially link cells of the same type, then cell-sorting occurs (Figure 1 D). Another feature we added from the literature is an implementation of individual cell migration: Polarized cells can migrate by pulling towards and pushing aside cells in front (Figure 1 A, E) (Delile, Doursat, et al., 2014; Delile, Herrmann, et al., 2017). We also implement functionality to use meshes for image-based modeling (Figure 1 F).

**Figure 1.**
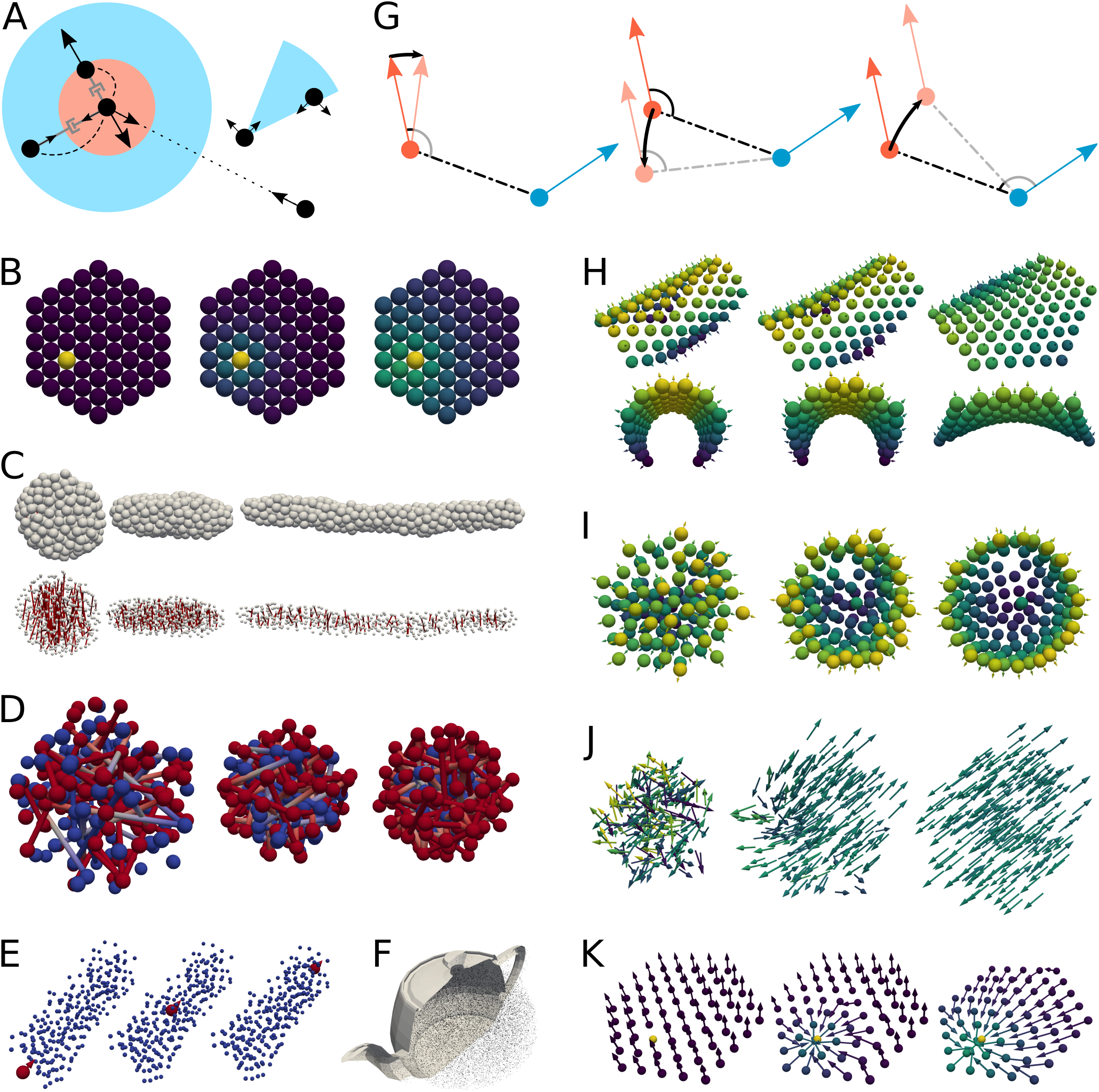
Spheroid Models Capture Many Cellular Behaviours. (A) Cells are described by their centers (black dots) surrounded by a repulsive core (red) and an attractive zone (blue). Cells within these ranges exert friction on each other (gray dashpots) and can exchange signals (dashed lines). More distant cells can be pulling towards each other by a contractile cellular protrusion (dotted line). Cells can migrate by pulling and pushing other cells (blue cone). (B) Exchange of signals leads to distributions that represent diffusive gradients. (C) Contractile protrusions can drive convergent extension or (D) cell sorting. (E) Migrating cell. (F) Image-based modeling, only half of the teapot mesh shown. (G) *U*_Epi_ at the red cell is minimized by the three depicted forces (black arrows). (H) Balancing these forces leads to layers with an elastic resistance to bending and (I) lets epithelial cells with initial polarities pointing radially outwards self-organize on a sphere (cut shown, simulated with friction on the background). (J) *U*_Pol_ aligns mesenchymal tissue polarity and (K) *U*_WNT_ reorients it. The latter is shown in a plane for clarity.

To this basic mesenchymal model we add two types of cell polarity – apical-basal polarity for epithelial cells and tissue polarity for mesenchymal cells. We represent these polarities by a spin-like unit vector 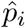 at each cell. For epithelial cells we introduce the potential

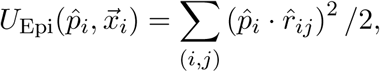

where the sum is over all pairs of neighbors in the same epithelium. This potential is minimal when each 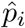 is orthogonal to all connections 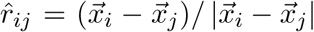 within the epithelium. The forces 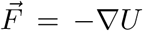 (Figure 1 G, see Supplemental Material for a derivation of the equations of motion) minimizing such a potential let cells self-organize into layers suitable to describe epithelial sheets (Figure 1 H, I).

Similarly, for mesenchymal cells’ tissue polarity we introduce the potential

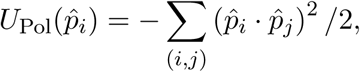

where the sum is over all pairs of mesenchymal neighbors. This potential is minimal when all polarities 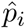 within the mesenchyme are parallel. It is therefore suitable to describe mesenchymal cells aligning due to tissue polarity (Figure 1 J). Diffusing signals like WNT are believed to act as an external influence to align tissue polarity (Yang and Mlodzik, 2015; Davey and Moens, 2017). Combining the ideas above we can simulate such behavior. The potential

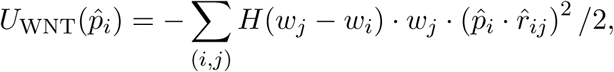

where *H* is the Heaviside function and *w* the signals concentration, orients polarities towards cells with higher concentration of *w* (Figure 1 K). The mesenchymal potentials *U*_Pol_ and *U*_WNT_ only induce torques on the polarities 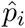 and leave the positions 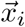 fixed.

We implemented all these spheroid models for GPUs (c.f. Supplemental Material for details), allowing us to simulate epithelial-mesenchymal interactions at large-scale. We demonstrate this by simulating how an epithelial Turing system induces branching (Figure 2 A) and how epithelial signals shape a tissue by controlling intercalation (Figure 2 B) (Menshykau et al., 2012). Moreover, our implementation for GPUs operates orders of magnitude faster than conventional implementations (Figure 2 C) (Marin-Riera et al., 2015; Mirams et al., 2013).

**Figure 2.**
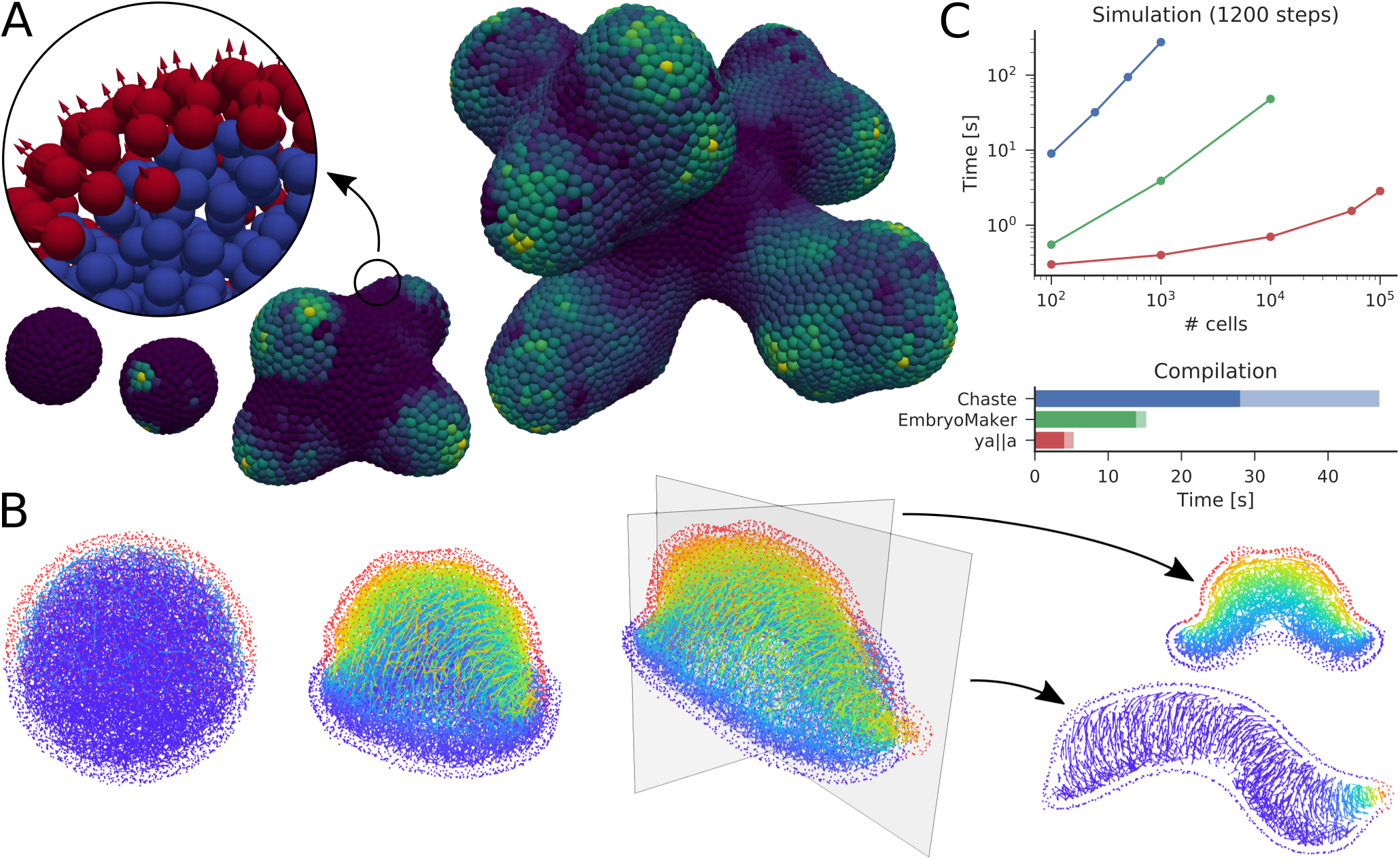
ya||a is Suitable for Large-scale Morphogenesis. (A) Branching driven by differential growth in the mesenchyme (blue) induced by Turing pattern on the epithelial surface (red). Coloring shows the concentration of the morphogen in the tissue in logarithmic scale, growing from 500 to roughly 40k cells in 4000 time steps. (B) Two epithelial signals shaping a tissue by controlling the distribution of protrusions, which induce intercalation. Cells intercalate along the gradient from top (shown in the series), except where this gradient is highest, there they intercalate normal to the second gradient from the tip (shown in the slice). The first rule folds the lower hemisphere into the upper hemisphere and the second rule elongates the structure. Slices show the two gradients in the final configuration. Around 12k cells, 500 time steps. (C) To compare computational performance we simulate cells with a limited interaction range, starting from a spherical, uncompressed, random distribution simulated until the forces were calculated 1200 times in Chaste (Mirams et al., 2013), EmbryoMaker (Marin-Riera et al., 2015) and ya||a. ya||a scales sub-linearly for small systems due to overhead. We use an Intel i7-4770 @ 3.40GHz with an NVidia GeForce GTX 1060 6GB. There would be a parallel implementation of Chaste scaling well for up to 32 CPU cores (Harvey et al., 2015).

## Discussion

Simulation frameworks for morphogenesis can be categorized into continuum models and agent-based models (ABMs) (Tanaka, 2015). Continuum models are based on descriptions of materials and describe bulk properties like viscosity or elasticity well at all timescales. However, it is difficult to interpret material properties like stiffness in terms of cellular behaviors and continuum models cannot be directly compared to cellular measurements like dispersion (Mogilner and Manhart, 2016). Furthermore, to numerically solve continuous models these must be discretized, usually using a mesh. Simulating growth and large deformations with meshes is challenging and computationally expensive (X. Chen and G. W. Brodland, 2008; Wittwer et al., 2016), however, often the bulk behavior on short timescales can be neglected. Then ABMs overcome the continuum model’s difficulties because they model cellular behaviors directly and agents provide a natural discretization.

To keep our simulations simple and fast we thus extended the spheroid model to include polarized cell behaviors, similarly to Hazelwood and Hancock (2013) for tissue polarity and similarly to Delile, Herrmann, et al. (2017) for epithelia. However, our model is inherently 3D and our potential for epithelia involves only pairwise interactions, which are easier to parallelize. Combining these novelties with previously proposed extensions for contractile protrusions (Palsson and Othmer, 2000; Belmonte et al., 2016) and individual cell migration (Delile, Herrmann, et al., 2017) makes the spheroid model ideal for large-scale simulations of 3D morphogenesis with mesenchyme and epithelium on equal footing.

For high performance at low costs we implemented these models into a GPU-based simulation framework. Current GPUs have thousands of cores, making them a cheap and comparably easy to program and use alternative to cluster computers. GPUs become increasingly popular for scientific computing (Nobile et al., 2017) their performance keeps improving. Even a cheap GPU allows us to calculate forces orders of magnitude faster than previous simulation packages (see Supplemental Material for hardware recommendations). Furthermore, outsourcing the heavy lifting to the GPU leaves the computer responsive enough for most other work during simulations.

We gain further performance over previous spheroid models (Mirams et al., 2013; Marin-Riera et al., 2015; Delile, Herrmann, et al., 2017) by implementing friction among neighbors, which lets deformations propagate through large systems and hence requires fewer time steps (Okuda et al., 2015).

We deliberately kept ya||a simple. Other packages support several kinds of models (Osborne et al., 2017) or provide sophisticated user interfaces (Marin-Riera et al., 2015; Delile, Herrmann, et al., 2017; Swat et al., 2012; Starruß et al., 2014). ya||a just works with spheroid models and relies on external programs for visualization. Thus the numerous tests and examples, including all models used to generate the figures, can be quickly understood and easily extended, because they are concise and plain CUDA/C++. This also leads to shorter compilation times, accelerating model development. The avoidance of dependencies additionally will make it easy to maintain ya||a in the future and the modular design will make it easy to integrate ya||a into larger pipelines, e.g. for parameter optimization.

While we are developing ya||a specifically for limb bud morphogenesis (Hopyan et al., 2011), similar cellular behaviors drive other developmental processes like tooth formation (Kim et al., 2017) or branching morphogenesis (Affolter et al., 2009) and computational modeling is becoming increasingly popular in developmental biology (Sharpe, 2017). Furthermore, epithelial-to-mesenchymal transitions are at the core of many cancers and cellular interactions are central to the immune system (Nagarsheth et al., 2017). The cell-centers 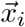 can even be re-interpreted as sub-cellular elements, then our model for epithelia could be used to describe cellular membranes (Milde et al., 2014). We therefore believe that ya||a will be very valuable to a wide community.

## Author Contributions

PG and MM developed ya||a under the supervision of JS. All authors wrote the manuscript.

## Acknowledgments

We thank Alexis Naveros, Giuseppe Bilotta, and Lorenzo Pistone for helping with CUDA, Stefanie Marti, Xavier Diego and Marco Musy for checking calculations, Xavier Diego for providing image-based meshes, Marco Musy for help with C++, Antoni Matyjaszkiewicz for tweaking the branching example, Marco Musy for careful reading of the manuscript, and James Osborne and the Chaste mailing list for helping with Chaste. This work was supported by the Spanish Ministry of Economy and Competitiveness, through ‘Centro de Excelencia Severo Ochoa 2013-2017’, SEV-2012-0208, by the Swiss National Foundation through Sinergia Grants CR23I3_156234 and CRSII3_141918, and by the European Research Council through SIMBIONT (project nr. 670555).

## Supplemental Methods

### Derivation of the Equations of Motion

Cells move with low inertia and high local friction, thus Newton’s equations 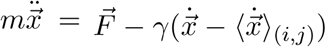 become 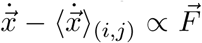 (Mao et al., 2013; Okuda et al., 2015). We usually use spherical forces like 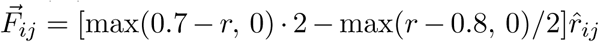 between cells 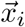 and 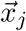 if 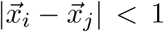, because they are simple to adapt. More complicated forces, like Lennard-Jones, require smaller time steps and do not qualitatively change behavior over long time scales. We simulate diffusion of a signaling molecule *w* at each cell *i* as

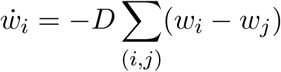

where the sum is over all neighbors of *i*. For protrusions we typically use constant forces and for proliferation we simply duplicate random cells, more sophisticated models can be easily implemented.

We describe polarities 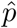 in spherical coordinates with *r* = 1, i.e.

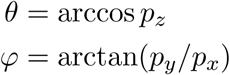

and conversely

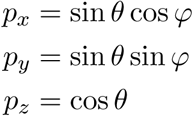

where 0 ≤ *θ* ≤ *π* and 0 ≤ *ϕ* < 2*π*. In these coordinates the scalar product is

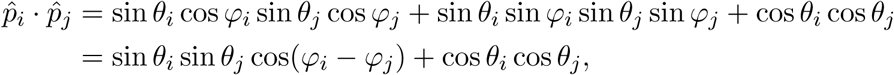

the gradient is

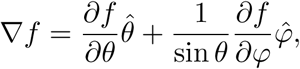

and the velocity is

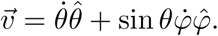

To find the equations of motion 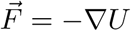 for 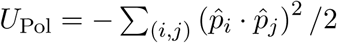 we therefore need to solve

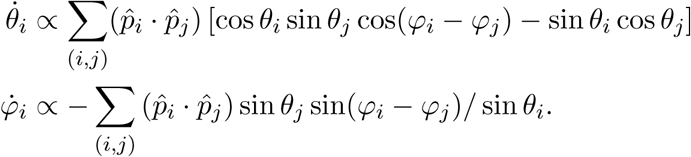

For 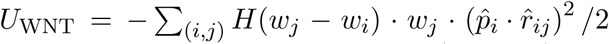 the second polarity 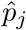 has to be replaced with 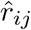 and a strength like *H*(*w*_*j*_ *-w*_*i*_) *· w*_*j*_ chosen.

Similarly, for 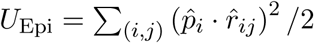 mainly the signs change, i.e.

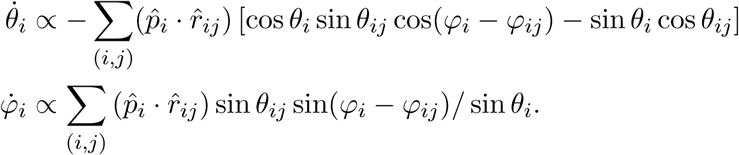

However, additionally contributions to 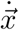 arise, because movement affects the angle between 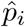 and 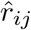 too. The contribution to 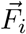 from the angle between 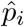 and *r*_*ij*_ is

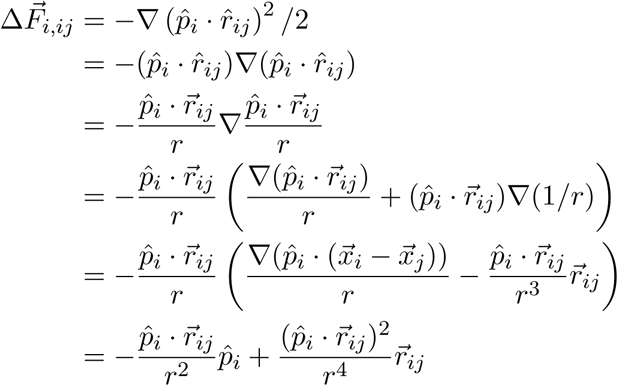

and similarly the contribution from the angle between 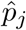 and *r*_*ji*_ is

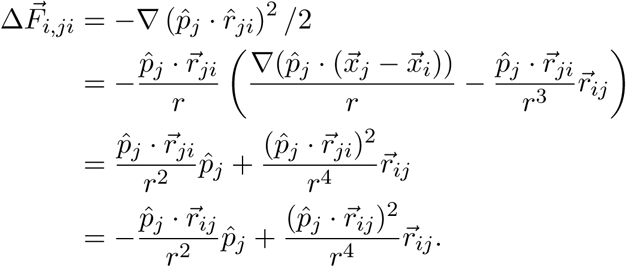

### Implementation

We approximate the equations of motion for 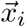 as 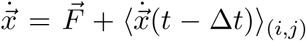. We solve the resulting ODE system using Heun’s method. Using a second order method allows taking approximately four times larger time steps without oscillations. The resulting time steps provide a suitable timescale for proliferation in our simulations, thus we do not use higher orders.

ya||a takes the definition of the right hand side in three parts, `Pairwise_interaction`, `Pairwise_friction`, and `Generic_forces`. The former two functions are called for each pair of neighbors and have to return a contribution to 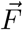 and a friction coefficient, respectively. The third, `Generic_forces`, can be used to compute further interactions that are not necessarily among neighbors, like the forces exerted by protrusions.

Most of the computational time is spent calculating the pairwise force and friction terms. For this we implemented two methods: `Tile_solver` calculates the interactions between each pair (Nyland et al., 2007) and `Grid_solver` uses a grid to identify possibly interacting pairs (Green, 2013). `Tile_solver` has less overhead and is thus faster for small systems, but calculation times grow with the square of the number of spheroids, while `Grid_solver` scales linearly. Our implementation is for Linux and macOS in CUDA/C++ and does not depend on additional libraries. We visualize the resulting Vtk files using Paraview (Ahrens et al., 2005).

Since the order of atomic operations is not deterministic, certain models produce different results in each run, due to accumulating numerical errors. While we could avoid many atomic operations we decided to embrace variation by seeding our random generators for each run, because we believe that biologically relevant models must be robust to this kind of noise.

We only use single (`float`) precision in our models. Single precision calculations run efficiently on the cheap consumer NVidia Geforce GPUs. Such GPUs can be installed in most computers, given enough power supply. Using various GPUs and profiling with nvvp we found that our calculations are memory bound, i.e. the execution speed scales with the memory bandwidth of the GPU.

While our early CUDA implementations were already comparably fast, they were held back by poor design. Initially, following *Numerical recipes in C* (Press et al., 1996), we used function pointers to pass the different interactions to our solvers. Using macros, and later templates, drastically improved performance. A second significant improvement was accumulating the forces in the local register, instead of writing each contribution directly into the GPU’s global RAM. The calculations are usually faster than writing the output, thus we use threads to generate output, while the GPU computes several steps.

We used Blender (Blender Online Community, 2017) to create a closed mesh for the teapot in Figure 1 F and Python packages for scientific computing (Waskom et al., 2017; McKinney, 2010; Hunter, 2007; Kluyver et al., 2016) to create the plot in Figure 2 C.

### Limitations

We have built ya||a for limb bud morphogenesis and we have not studied unrelated morphogenetic processes, like apical constriction or planar cell polarity. However, ya||a is very flexible and developing corresponding models would be straight forward.

Similarly, the solvers we require are based on the overlapping spheres model, i.e. all cells within a certain radius are potentially interacting neighbours. Such a model is not ideal for processes with cells deforming drastically before changing neighbors (Pathmanathan et al., 2009; Osborne et al., 2017). While alternative interaction models can be implemented on top of the provided solvers, the required code might be more complex and the resulting simulations might be slower than dedicated solvers.

Running ya||a requires a graphics card from NVidia, since it is implemented for NVidia’s parallel computing platform CUDA. CUDA supports many generic programming features and includes valuable resources like a visual profiler, making CUDA easier to program than more portable platforms. This platform choice affects mainly laptops as other computers can be upgraded at low cost.

